# Structural and Functional Analyses of Hub MicroRNAs in an Integrated Gene Regulatory Network of *Arabidopsis*

**DOI:** 10.1101/2020.06.10.145185

**Authors:** Zhaoxu Gao, Jun Li, Li Li, Yanzhi Yang, Jian Li, Chunxiang Fu, Danmeng Zhu, Hang He, Huaqing Cai, Lei Li

**Author notes:** Corresponding author. (Li L). Equal contribution (Zhaoxu Gao, Jun Li).

## Abstract

MicroRNAs (miRNAs) are trans-acting small regulatory RNAs that work coordinately with transcription factors (TFs) to shape the repertoires of cellular mRNA available for translation. Despite our growing knowledge of individual plant miRNAs, their global roles in gene regulatory networks remain mostly unassessed. Based on interactions reanalyzed from public databases and curated from the literature, we reconstructed an integrated miRNA network in *Arabidopsis* that includes 66 core TFs, 318 miRNAs, and 1712 downstream genes. We found that miRNAs occupy distinct niches and enrich miRNA-containing feed-forward loops (FFLs), particularly those in which the miRNAs are intermediate nodes. Further analyses revealed that miRNA-containing FFLs coordinate TFs located in different hierarchical layers and that intertwined miRNA-containing FFLs are associated with party and date miRNA hubs. Using the date hub *MIR858A* as an example, we performed detailed molecular and genetic analyses of three interconnected miRNA-containing FFLs. These analyses revealed individual functions of the selected miRNA-containing FFLs and elucidated how the date hub miRNA fulfills multiple regulatory roles. Collectively, our findings highlighted the prevalence and importance of miRNA-containing FFLs to provide new insights into the design principles and control logic of miRNA regulatory networks governing gene expression programs in plants.

## Introduction

Gene expression programs are fundamental to organism integrity and function. These programs are encoded and decoded by regulatory networks based on interactions between trans-acting factors and cis-regulatory elements. Transcription factors (TFs) and miRNAs are two primary classes of regulators with defined target specificity as well as the ability to substantially impact the transcriptome [1]. Arising from primary transcripts, pre-miRNAs with intramolecular stem-loop secondary structures are further processed by evolutionarily conserved cellular machinery (DICER-like complexes in plants) to yield mature miRNAs [2]. Similar to TFs, miRNAs act in trans and recognize their targets through short sequence motifs termed miRNA binding sites (MBSs) [1]. However, miRNAs function mainly at the post-transcriptional level to guide target cleavage or translational inhibition, thereby complementing TF-based transcriptional control of mRNA synthesis [1, 2].

Given the crucial role of miRNAs in gene regulation, integrated networks including both transcriptional and post-transcriptional regulation are necessary to provide a comprehensive view of gene expression programs and to elucidate their design principles [3, 4]. In animals, many studies have reported reconstruction of such integrated networks, efforts that have helped to reveal the interconnected global relationship between miRNAs and other regulatory agents [4-6]. Although mutant and transgenic analyses have associated specific miRNAs with particular biological processes in plants, systematic efforts aimed at elucidating plant miRNA networks are largely lacking [7, 8]. Consequently, the joint action of miRNAs and TFs in modulating the transcriptome, and thus the plant phenome, has not been fully described.

Network motifs are subgraphs that occur more often in real-world networks than in random connections [9, 10]. Conserved network motifs present in diverse species represent evolutionarily successful mechanisms for regulating gene expression. In particular, the three-node-three-edge feed-forward loop (FFL), which contains a direct path between the input node and the output node and an indirect path via an intermediate node, is a prominent and versatile network motif found in transcription networks of both prokaryotic and eukaryotic origins [9, 11-13]. In plants, a number of FFLs that play diverse regulatory roles in multicellular development and in response to biotic and abiotic stresses have been experimentally studied [14-16].

Studies in animals combining bioinformatics and experimental analyses have revealed that miRNAs are often found within network motifs containing TFs [17, 18]. Animal miRNAs require as little as 6-8 nucleotides for effective targeting [19, 20], which entails enormous connectivity among the targets, such that various RNA species may become “competing endogenous RNAs” and react with each other through the same MBS [21]. By contrast, miRNA-target pairs in plants typically display near-perfect complementarity [22, 23], implying that the target spectra of plant miRNAs are narrower than that in animals. How this difference dictates architectural and topological characteristics of plant miRNA networks remains to be evaluated. Moreover, most plant miRNA loci are independent transcriptional units subject to RNA polymerase II–based regulation [24, 25]. The large numbers of TFs encoded in plant genomes potentially form a myriad of regulatory interactions with miRNA loci, a relationship which also remains to be explored in plants [8].

Hubs in the gene regulatory networks represent a small proportion of nodes that exhibit maximal information exchange with other nodes. Network hubs include party and date hubs [10, 26]. A party hub has all its downstream nodes located within a module, and these nodes act together to regulate a biological process. By contrast, a date hub has many interactions across multiple modules that function in different contexts [10, 26]. To date, plant miRNAs have not been systematically analyzed for their association with network hubs. Further mapping and analysis on a genome-scale is expected to provide important mechanistic insights into the miRNA gene networks that underpin plant development and responses to environmental challenges [8, 10, 26-28].

In the current study, we reconstructed a comprehensive miRNA regulatory network in *Arabidopsis* by integrating three distinct types of interactions: miRNA-target interactions (MTIs), TF-miRNA interactions (TMIs), and TF-target interactions (TTIs). Examination of this directed network, which consisted of 66 core TFs, 318 miRNAs, and 1712 miRNA target genes, revealed that miRNAs occupy unique niches and enhance the formation of miRNA-containing FFLs. Global analysis coupled with comprehensive experimental characterization of selected FFLs related to the date hub *MIR858A* demonstrated that miRNAs have profound effects on the hierarchical organization and control logic of gene expression programs. Collectively, our results provide an example of using combined systems and molecular biology approaches to elucidate the structural and functional roles of plant miRNAs in the context of regulatory networks.

## Results

### Reconstruction of an integrated miRNA network in *Arabidopsis*

As previously recommended [29], we incorporated multiple sources to comprehensively identify high confidence MTIs in *Arabidopsis* (**Figure 1**A). Starting from 428 annotated miRNAs, we generated four combined outputs from the psRobot [30] and psRNATarget [31] programs using two stringency levels (Figure S1A). The dataset with the most optimal tradeoff between recovery of a high portion of the canonical targets and minimization of the overall number of predictions was retained (Figure S1B). Together with 111 validated MTIs not retrieved by the above pipeline, a final set of 2823 MTIs was compiled (Figure S1C), covering 318 (74.5%) miRNAs and 2008 target genes. Mapping the MBS against the gene structure revealed a distribution slightly biased towards the 5’ untranslated region (UTR) and the distal end of the coding region (Figure S1D). Of the 1717 *Arabidopsis* TF genes, 203 from 35 families were included in the MTIs (Figure S2A), a frequency significantly higher than the genome average (Figure S2B). Gene Ontology (GO) analysis further revealed that target genes were enriched for terms related to gene expression, response to abiotic/biotic stimulus, and development (Figure S2C).

**Figure 1.**
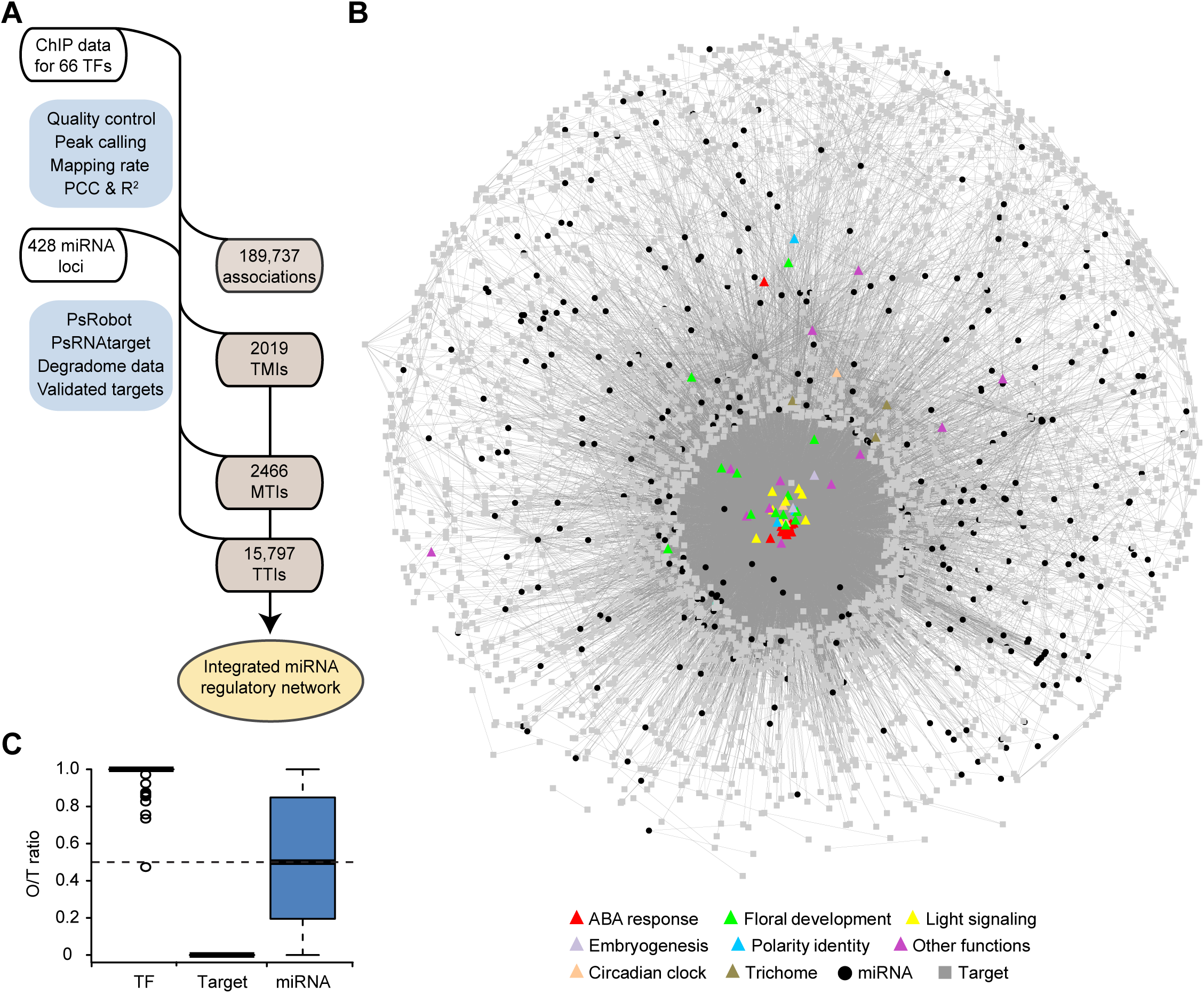
Reconstruction of an integrated miRNA network in *Arabidopsis*. **A**. Diagram illustrating the workflow for reconstructing a miRNA regulatory network in *Arabidopsis*. MTIs were computationally predicted and curated from the literature. TMIs and TTIs were based on uniformly reprocessed ChIP data. **B**. Visualization of the reconstructed network in the edge-weighted spring embedded layout format of Cytoscape. Core TFs are shown as triangles and are colored according to their annotated functions. MiRNAs and other genes are depicted as black cycles and grey squares, respectively. All edges are represented as grey lines. **C**. The average O/T ratio for different network components. The out-degrees for the core TFs, miRNAs, and target genes were individually calculated and divided by their total degree. The dashed line represents the theoretical O/T ratio (0.5) at the network level.

To map TTIs and TMIs, we exploited published whole genome chromatin immunoprecipitation (ChIP) data. After quality control and uniform processing, we identified 339,875 binding peaks for 66 TFs (hereafter referred to as core TFs; Figure S3A; Table S1). The binding peaks were predominantly located in the non-centromeric chromosome space (Figure S3B). In relation to gene structure, most binding peaks were found in intergenic regions, followed by exons, introns, 5’ UTRs, and 3’ UTRs (Figure S3C). For protein coding genes, the peaks were concentrated around the transcription start site (TSS; Figure S4A). For miRNA loci, the peaks were predominantly located at a region approximately 200 bp upstream of the first nucleotide of the annotated pre-miRNA (Figure S4B), an observation consistent with previous analysis of RNA polymerase II binding profiles [24, 25]. By defining a binding window to link a peak to a gene (Figure S4), we identified 189,737 associations between the core TFs and 26,023 downstream genes, including 2019 TMIs for 263 miRNAs and 15,797 TTIs for 1712 miRNA targets (Figure 1A). Integrating the MTIs, TMIs, and TTIs, we reconstructed a miRNA regulatory network consisting of 2096 nodes (genes) and 20,282 directed edges (interactions) (Figure 1B).

### MiRNA nodes preferentially form FFLs

To describe the general properties of the miRNA nodes, we calculated degree distribution, which refers to the number of direct interactions through which a node is linked to other nodes. We found that the overall degree distribution follows a power-law, with a small number of nodes (hubs) having extremely high connectivity (Figure S5A). Since the network is directed, we further considered the in-degree (regulated) and out-degree (regulating) of the nodes and found the in-degree distribution also follows a power-law (Figure S5B and S6A). The path length is rather short, with the majority of paths having one to three steps (Figure S5C). The mean connectivity for the miRNA nodes is 14.1, greater than the network average of 9.68. Removal of the miRNA nodes did not significantly change the network topology with regards to clustering coefficients and closeness centrality but did reduce the betweenness centrality (Figure S5D and S5E). Moreover, the miRNA nodes possess more balanced out-degree and in-degree, with an average out-degree over total degree (O/T) ratio near 0.5 (Figure 1C). Taken together, these observations suggest that the miRNA nodes are more connected than other nodes.

We identified several enriched motifs in the network (Figure S6B). FFL is an important network motif in transcription networks whose dynamic properties have been comprehensively analyzed [11, 13]. In the reconstructed network, a total of 19,615 FFLs were identified, significantly enriched compared to the permutated networks (Z score > 2.0), while the isomeric feedback loops were underrepresented (**Figure 2**A). Although miRNAs accounted for only 15.2% of the nodes, a significantly higher portion (35.6%, *P* < 0.001) was contained in the FFLs (Figure 2B). Moreover, we found that the conserved miRNAs, which could be found in species other than *Arabidopsis*, are included in the FFLs in an even higher proportion (Figure 2C). Among the 6976 miRNA-containing FFLs, 5334 (76.5%) position the miRNA as the intermediate node (Figure 2D). FFLs containing conserved miRNAs have the same configurations regarding the position of the miRNA (Figure 2E and 2F). Together, these results indicate that the miRNA nodes occupy distinct niches in the network to preferentially form FFLs.

**Figure 2.**
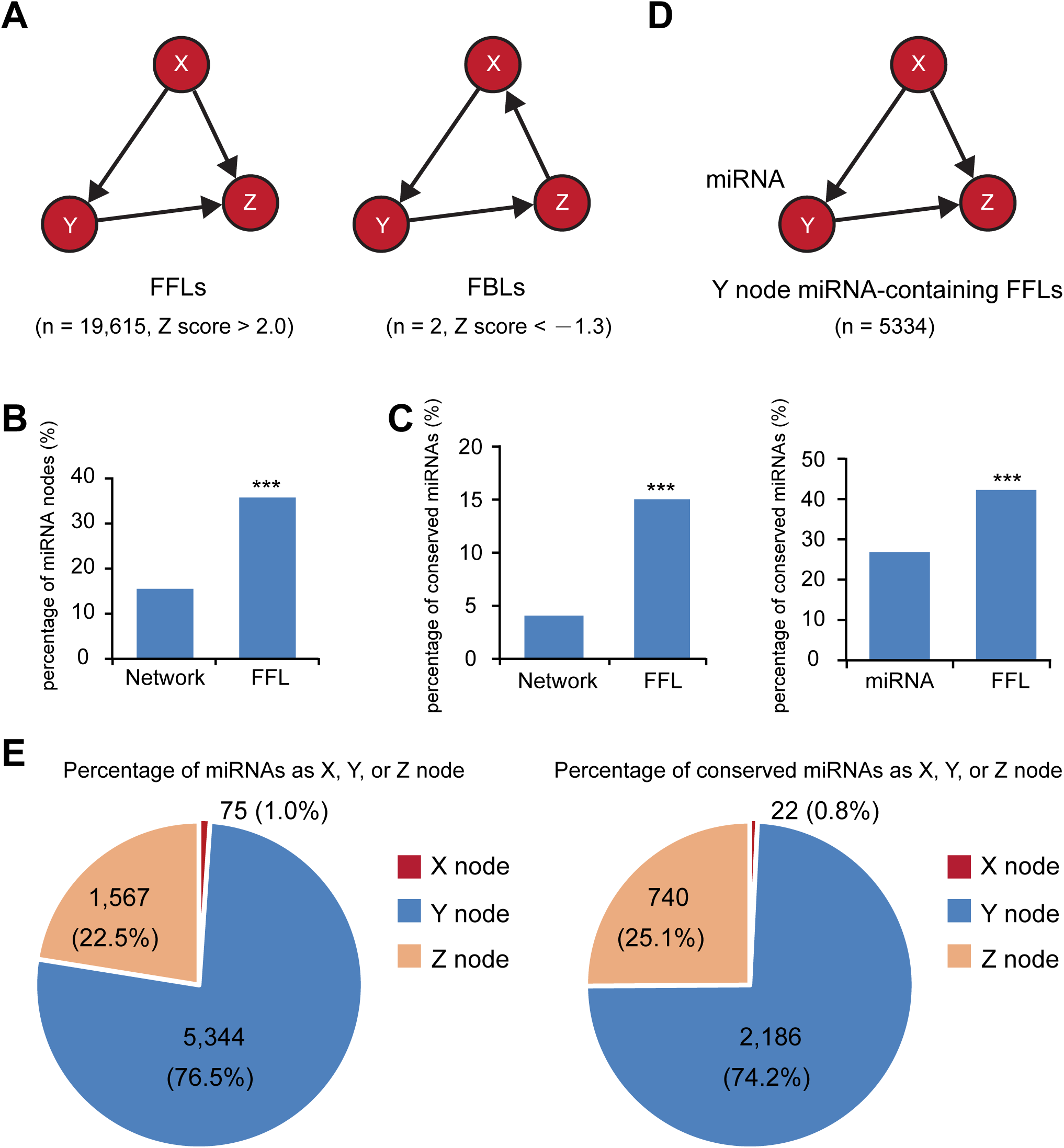
MiRNA-containing FFLs are enriched in the reconstructed network. **A**. Diagram of the three-node-three-edge FFL and FBL subgraphs that differ only in the direction of the direct path between the X and Z nodes. Z scores for the frequency of the two subgraphs in the reconstructed network are shown. **B**. Comparison of the proportion of miRNA nodes in the network and in the FFLs. *** indicates *P* < 0.001 by chi-square test. **C**. Analysis of FFLs containing conserved miRNAs. The bar graph on the left shows the proportion of conserved miRNAs in the network and in the FFLs. The graph on the right shows the proportion of conserved miRNAs in total network miRNAs and the proportion of FFLs containing conserved miRNAs in total miRNA-containing FFLs. **D**. Diagram of a miRNA-containing FFL in which the Y node is a miRNA. **E**. Pie graphs showing the proportion of FFLs in which the miRNAs (left) and conserved miRNAs (right) are positioned as the X, Y, or Z node.

### Date hub miRNAs link multiple intertwined FFLs

Based on clustering coefficient and degree level, we identified 26 miRNA hubs in the reconstructed network (**Figure 3A**). These hubs were further classified into ten putative party hubs and 16 date hubs, based on whether or not the target genes are associated with similar GO terms (Figure 3A). For example, *MIR408* is a representative party hub, as its target genes encode predominantly cuproproteins and are associated with highly similar GO terms. Conversely, *MIR858A* is a typical date hub that targets multiple *MYB* family members associated with dissimilar GO terms. Isolated subgraphs concerning *MIR408* and *MIR858A* revealed top- and bottom-heavy structures, respectively (Figure 3B), resembling multi-input and multi-output FFLs that might have derived from miRNA-containing FFLs through topological generalization [17]. For example, *MIR858A* underpins 91 FFLs that connect with different output nodes, while *MIR408* has only two FFLs (Figure 3B). Quantification of FFLs revealed that date hubs indeed exhibited significantly more FFLs than the party hubs that were compatible with topological generalization of the output node (Figure 3C).

**Figure 3.**
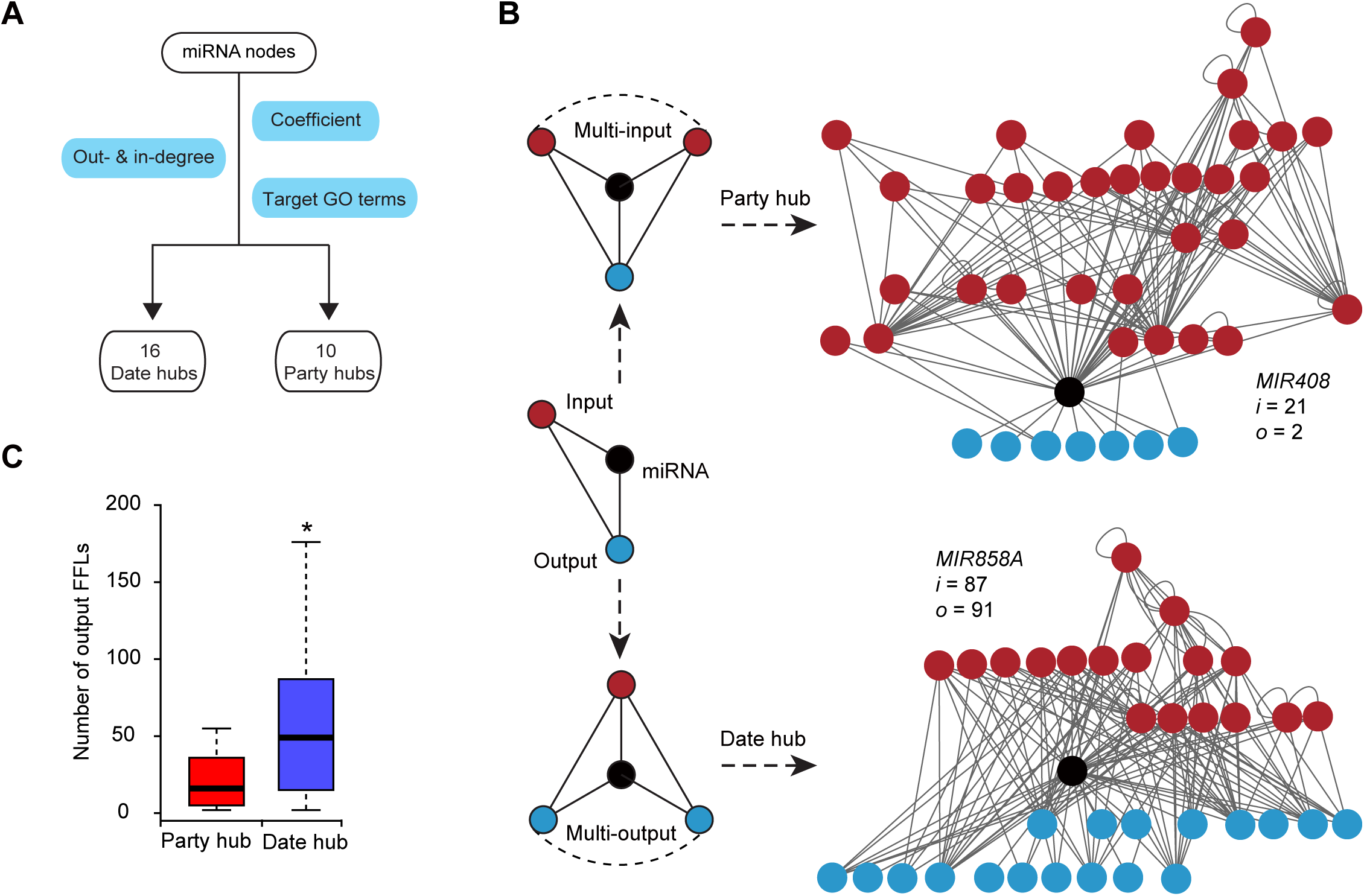
Identification of date hubs and party hubs from the miRNA network. **A**. Flowchart showing the process for identifying various hub miRNAs. The in hubs and out hubs were classified based on their in-degree, out-degree, and clustering coefficient. Party and date hubs were further distinguished by manual inspection of similarity or dissimilarity of the GO terms associated with their target genes. **B**. Scheme of party and date hub formation through topological generalization of miRNA-containing FFLs (black cycle). The input-heavy and output-heavy structures of the party (e.g., *MIR408*) and date hubs (e.g., *MIR858A*) result from preferential topological generalization on the input (red cycle) and the output node (blue cycle) of a miRNA-containing FFL, respectively. The numbers “*i*” and “*o*” represent degree of deduced generation for the input and output node, respectively. **C**. Box plot showing the number of output FFLs by topological generation of the putative party and date miRNA hubs. *, *P* < 0.05 by chi-square test.

To study the date hubs, we used *MIR858A* as an example and sought to validate its targeting of multiple *MYB* genes. Available degradome sequencing data provided evidence for at least ten miR858-*MYB* interactions (**Figure 4**A). We also performed a 5’ RNA ligase mediated rapid amplification of complementary DNA ends (5’ RLM-RACE) assay (Figure 4B). Similar to previous reports [32], we found that the cloned 5’ ends are located at or near the predicted cleavage sites (Figure 4B). To validate the miR858-*MYBL2* interaction with the negative 5’ RLM-RACE result, we modified a previously reported REN/LUC dual-luciferase system [33], in which the *MYBL2* coding region was fused with *LUC*. We also generated *MYBL2*^*mMBS*^*-LUC* by substituting nucleotides in the miR858 MBS while keeping the amino acid sequence intact (Figure 4C). The dual-reporter constructs were used to transiently transform tobacco protoplasts, which revealed that normalized LUC activity from *MYBL2*^*mMBS*^*-LUC* was significantly higher than that from *MYBL2-LUC* when co-transformed with miR858 (*P* < 0.05) (Figure 4D), indicating that miR858 is able to attenuate MYBL2 accumulation.

**Figure 4.**
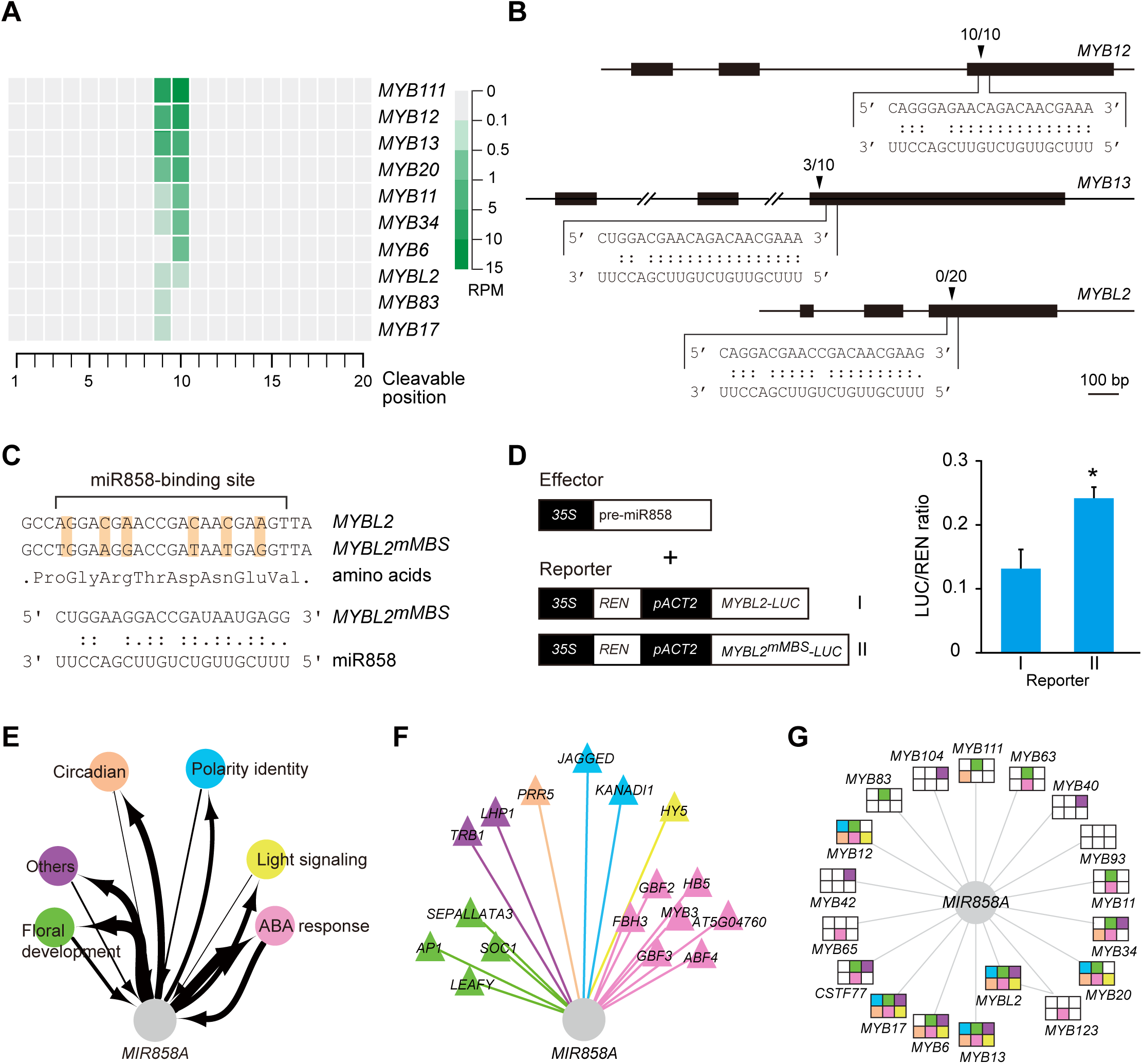
Experimental validation of *MIR858A* as a date hub. **A**. Degradome sequencing data support miR858 targeting of multiple *MYB* family members. The 20 possible miR858 guided cleavage positions in the predicted miR858 binding sites of 10 *MYB* transcripts are aligned. Frequency of the sequenced ends mapped to the middle three positions is displayed using the color key shown at the bottom. **B**. Analysis of miR858 targeting of three *MYB* genes by 5’ RLM-RACE. Gene structures are shown at the top. Base pairing between the MBSs and miR858 is shown on the bottom, with “.” indicating G:U wobble pairing. Triangles mark the cleavage sites along with the frequency of the corresponding clones in the RACE assay. **C**. *MYBL2*^*mMBS*^*-LUC* was created by substituting the shaded nucleotides in the MBS but maintaining the amino acid sequence. **D**. *35S:pre-miR858a* (effector) and the *MYBL2-LUC* or *MYBL2*^*mMBS*^*-LUC* reporter were used to transiently co-transform tobacco protoplasts. Data are means ± SD (n = 3) of the calculated LUC/REN chemiluminescence ratio. *, *P* < 0.05 by one way ANOVA test. **E**. Diagram illustrating participation of *MIR858A* in six of the eight sub-networks, which were extracted from the pan-network based on the functions of the core TFs. Arrows depict TMIs and MTIs, with the line thickness indicating node degree. **F**. Detailed wiring pattern among TF nodes neighboring *MIR858A*. TMIs are color-coded based on assignment of the given TFs to the sub-networks. **G**. Analysis of MTIs between *MIR858A* and the 18 target genes. Assignment of the target genes to the six sub-networks is represented by shaded squares. The color schemes used in panel F and G are the same as in E.

To further analyze the hub properties of *MIR858A*, we extracted eight modularized sub-networks based on their core TFs (Figure S7A and S7B). In accordance with the functions of the TFs, the individual sub-networks found to relate to ABA response, floral development, light signaling, circadian rhythms, polarity identity, trichome, embryogenesis, and other functions (Figure S7A and S7B). The clustering coefficient of the TF nodes was more constant in the sub-networks than at the individual TF level (Figure S7C), attesting to the modularity of the isolated sub-networks. We found that the *MIR858A* node is connected to six of the eight sub-networks through TMIs and MTIs (Figures 4E to 4G). These results confirmed that *MIR858A* bridges multiple functional modules defined by TFs, thereby facilitating crosstalk among diverse biological processes to constitute a date hub.

### Date hubs fulfill pleiotropic effects

To functionally characterize *MIR858A*, we dissected out three interconnected FFLs (**Figure 5A**) from our network. This selection was based on the *HY5-MIR858A-MYBL2* FFL, for which the molecular interactions were supported in the literature [34, 35]. FFL is either coherent or incoherent depending on whether the direct and indirect paths have the same net regulatory effects [11]. After performing additional experiments to elucidate the signs of the *HY5-MIR858A* (Figure S8) and *HY5-MYBL2* edges (Figure S9), we concluded that *HY5-MIR858A-MYBL2* is a coherent FFL, with the TTI and TMI emanating from the input TF *HY5* displaying opposite signs of regulation (Figure S8A). Direct visual (Figure 5B) and chemical quantification (Figure 5C) of mutants disrupting each node revealed that this FFL plays a role in regulating light induced anthocyanin accumulation in seedlings.

**Figure 5.**
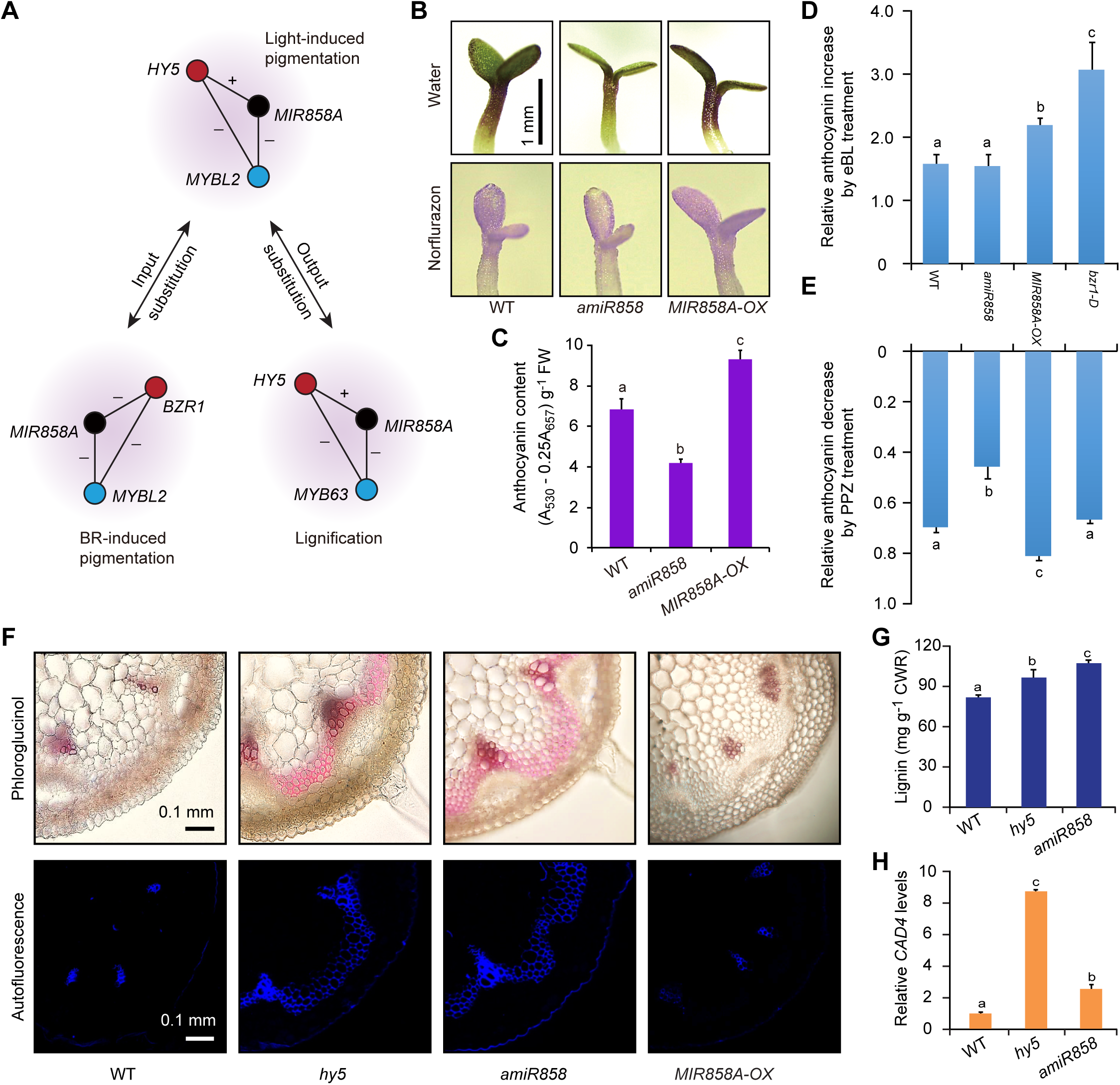
The date hub *MIR858A* fulfills pleiotropic effects through intertwined FFLs. **A**. The intertwined interactions centered on *MIR858A* (black) are dissected into three FFLs, whereby substitution of the input (red) and output nodes (blue) of the *HY5-MIR858A-MYBL2* FFL generates the *BZR1-MIR858A-MYBL2* and *HY5-MIR858A-MYB63* FFLs, respectively. +, positive regulation; -, negative regulation. **B**. Comparison of anthocyanin accumulation in seedlings grown for 3 days on media in the light, with water and norflurazon treatments. Bar, 1 mm. **C**. Quantitative measurement of anthocyanins in seedlings of the indicated genotypes. **D**. Quantification of eBL-induced anthocyanin accumulation. Seedlings were treated with ethanol (mock) or 0.1 μM eBL. The relative increase in anthocyanin level ([eBL-ethanol]/ethanol) was determined for the four indicated genotypes. **E**. Quantification of PPZ induced anthocyanin suppression. Seedlings were treated with DMSO (mock) and 0.1 μM PPZ. The relative decrease in anthocyanin level ([DMSO-PPZ]/DMSO) was determined. **F**. Examination of lignin accumulation in the four indicated genotypes. Early-stage inflorescence stems were cross-sectioned and examined by phloroglucinol staining and UV-excited autofluorescence. Bars, 0.1 mm. **G**. Quantification of lignin level in young stems of the indicated genotypes. **H**. qPCR analysis of transcript level of the lignin biosynthesis gene *CAD4* in early-stage stems. Data are all means ± SD (n = 3). Different letters denote groups with significant differences (one way ANOVA test, *P* < 0.05).

Substituting the input node *HY5* with *BRASSINAZOLE-RESISTANT 1* (*BZR1*), a key TF in Brassinosteroid (BR) signaling, produced the *BZR1-MIR858A-MYBL2* FFL (Figure 5A). In addition to transcript analysis, *pMIR858A:GUS* and *pMYBL2:GUS* reporter lines displayed weakened GUS activity after exogenous treatment with 2, 4-epicastasterone (eBL), a synthetic BR analog (Figure S10A and S10B). Conversely, exogenous treatment with propiconazole (PPZ), a BR biosynthesis inhibitor, resulted in increased miR858 and *MYBL2* transcript levels (Figure S10C and S10D). These results indicate that BR represses *MIR858A* and *MYBL2* expression and together with the fact that miR858 negatively regulates *MYBL2*, demonstrate that *BZR1-MIR858A-MYBL2* is an incoherent FFL (Figure 5A).

Consistent with previous reports [36, 37], we found that eBL induces anthocyanin accumulation (Figure 5D; Figure S10E). Exogenous application of PPZ resulted in reduced pigmentation (Figure 5E; Figure S10E). In the *bzr1-1D* mutant with constitutive BR signaling [38], the effect of exogenous eBL on pigmentation was enhanced (Figure 5D). *MIR858A-OX* was able to phenocopy *bzr1-1D* with regards to eBL-induced anthocyanin accumulation (Figure 5D). Conversely, suppression of pigmentation by PPZ treatment was alleviated in *amiR858* (Figure 5E). Based on these results, we concluded that miR858 is a positive regulator of BR-induced anthocyanin biosynthesis. Thus, by sharing the *MIR858A-MYBL2* output circuit, light and BR converge to regulate seedling pigmentation.

*HY5-MIR858A-MYB63* is another coherent FFL in which the output node *MYBL2* is substituted with a different MYB family member (Figure 5A; Figure S10F). The known hyper-lignification phenotype of *MYB63-OX* [39] prompted us to investigate whether the *HY5-MIR858A* circuit is involved in lignin formation. Cytochemical staining and analysis of lignin autofluorescence of various mutants revealed that *amiR858* and *hy5* exhibited increased stem lignin deposition compared to the wild type (Figure 5F), a phenotype consistent with that of *MYB63-OX* [39]. Quantification of lignin content (Figure 5G) and transcript analysis of lignin biosynthesis genes (Figure 5H) confirmed the lignin hyper-accumulation phenotype of *amiR858* and *hy5*. Thus, the *HY5-MIR858A-MYB63* FFL is critical for suppression of *MYB63* expression and maintenance of proper lignin levels during inflorescence stem development. Collectively, our results demonstrate that *MIR858A* participates in modulating multiple *MYB* family members involved in different processes and thus constitutes a functional date hub fulfilling multiple regulatory effects through interconnected FFLs.

### FFLs coordinate TFs from different hierarchical layers

The intricate connection between date hub miRNAs and TFs prompted us to isolate a TF-miRNA core-network consisting of 249 TFs and 275 miRNAs along with 5177 edges (**Figure 6A**). Based on clustering coefficient and degree level, we divided the core-network into top, middle, and bottom layers, which include 48 (9.2%), 18 (3.4%), and 458 (87.4%) nodes, respectively (Figure 6A). To test whether this hierarchical organization is associated with gene regulation, we compared transcriptomic profiles of all TF genes between the wild type, *ago1*, which is defective in *ARGONAUTE 1* required for miRNA action, and *rdr6*, which is defective in *RNA-DEPENDENT RNA POLYMERASE 6* involved in siRNA biogenesis. We observed that the core TFs, which were mainly located in the top layer (*P* < 0.05), were not significantly influenced in the mutants (Figures 6A and 6B; Figure S11A). Of the 249 TFs in the core-network, 193 (77.5%) were miRNA targets (Figure S11B). Consistent with this finding, the average expression level of these TFs was significantly higher than the non-targets in the *ago1* mutant but not in *rdr6* (*P* < 0.01) (Figure 6C). Regarding the hierarchical layers, we found that the average expression level of the bottom-layer genes was significantly higher than that of the top-layer in *ago1* but not *rdr6* (Figure 6D). Together, these results indicate that miRNAs specifically influence expression of the target TFs as well as TFs in the bottom hierarchical layer.

**Figure 6.**
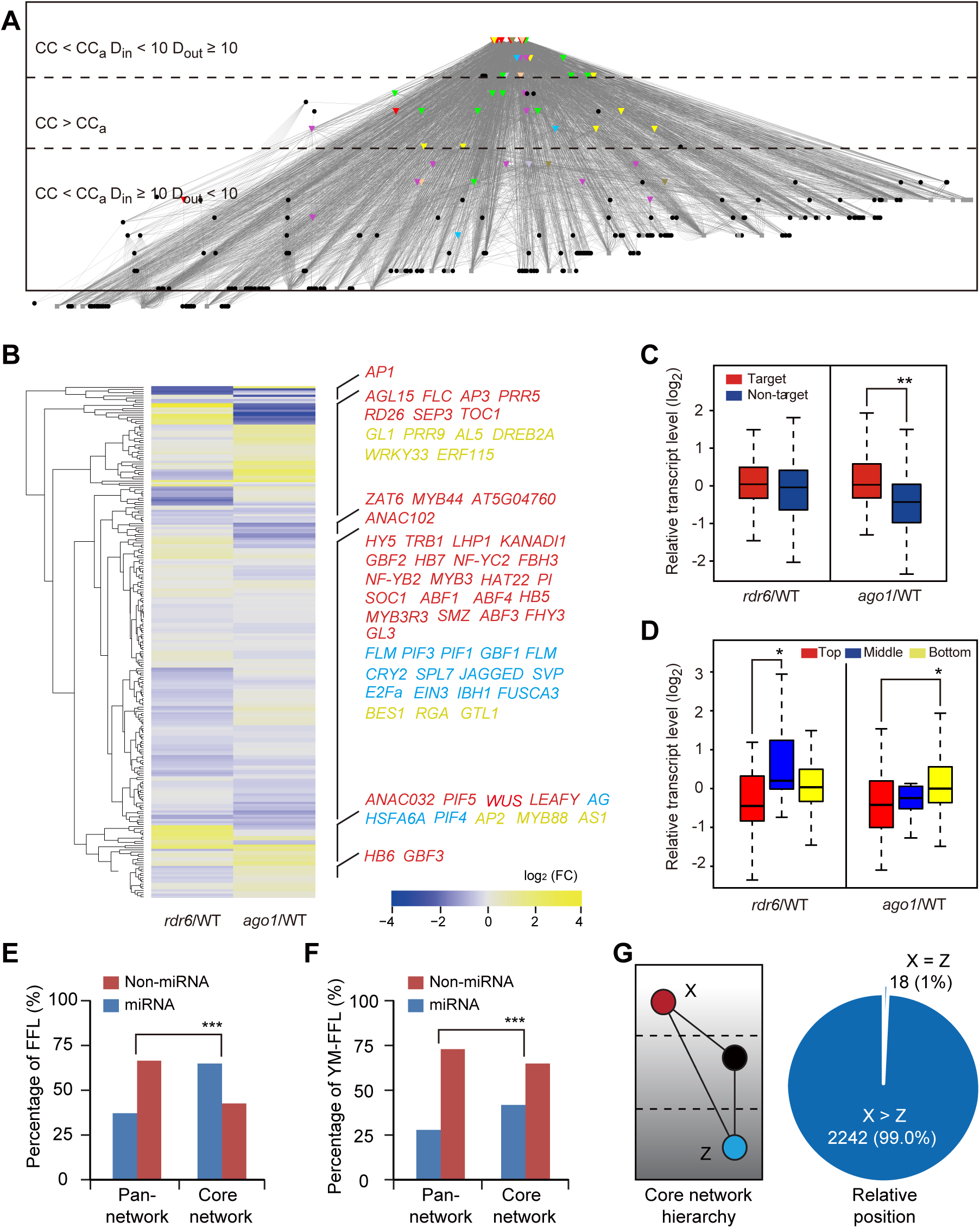
Topological and regulatory hierarchy of the TF-miRNA core network. **A**. A TF-miRNA core network in which the core TFs are depicted as colored triangles, miRNAs as black circles, and other TFs as grey squares. The top, middle, and bottom hierarchical layers were divided based on their node clustering coefficients and degree distributions. CC, clustering coefficient; CC_a_, average clustering coefficient of the pan-network; D_in_, number of in-degree; D_out_, number of out-degree. **B**. Transcriptomic influence of small RNA pathways on the core network. The heatmap shows the relative expression levels of the 249 TFs in *rdr6* and *ago1* compared to the wild type. The positions of the core TFs are indicated on the right. **C**. Boxplot showing expression of the 193 miRNA targets and 56 non-targets in *rdr6* and *ago1* in comparison to the wild type. **, *P* < 0.01 by chi-square test. **D**. Boxplot analysis of relative expression of the TF genes that were divided by their hierarchical layers. *, *P* < 0.05 by chi-square test. **E**. The proportions of FFLs with or without a miRNA are significantly different in the core network in comparison to that in the pan-network (***, *P* < 0.001 by chi-square test). **F**. The proportion of FFLs with an miRNA as the intermediate node is significantly higher in the core network than in the pan-network (*P* < 0.001). **G**. Relating miRNA-containing FFLs to the hierarchy of the core-network. The hierarchical layer of the nodes was quantified, whereby X > Z if X is present in a layer higher than Z (left). The pie graph (right) shows that X is essentially always in a higher layer than Z for miRNA-containing FFLs.

Of the 6296 FFLs in the core-network, the majority (3836 or 61%) were miRNA-containing, which represents a significant enrichment compared to the pan-network (Figure 6E). Further, miRNAs were positioned as the intermediate (Y) nodes in 2256 (35.8%) of the FFLs, also significantly higher than observed in the pan-network (27.2%) (Figure 6F). We further found that the X (input) node was almost always in a hierarchical layer higher than the Z (output) node in miRNA-containing FFLs (Figure 6G). To relate this structural feature to biological relevance, we employed the exemplar *HY5-MIR858A-MYBL2* FFL for further analysis. Anthocyanin biosynthesis is divided into early and late stages [40]. While *MYBL2* interferes with formation of the transcription complexes that activate late-stage genes (**Figure 7A**) [40, 41], we found that HY5 binds especially to the promoters of early-stage genes (Figure 7A), based on the global ChIP data [7]. Thus, the positioning of the input and output nodes along the anthocyanin pathway is consistent with the hierarchy of the FFL in the core network, with the three nodes locating to the top, middle, and bottom layers (Figure 7B).

**Figure 7.**
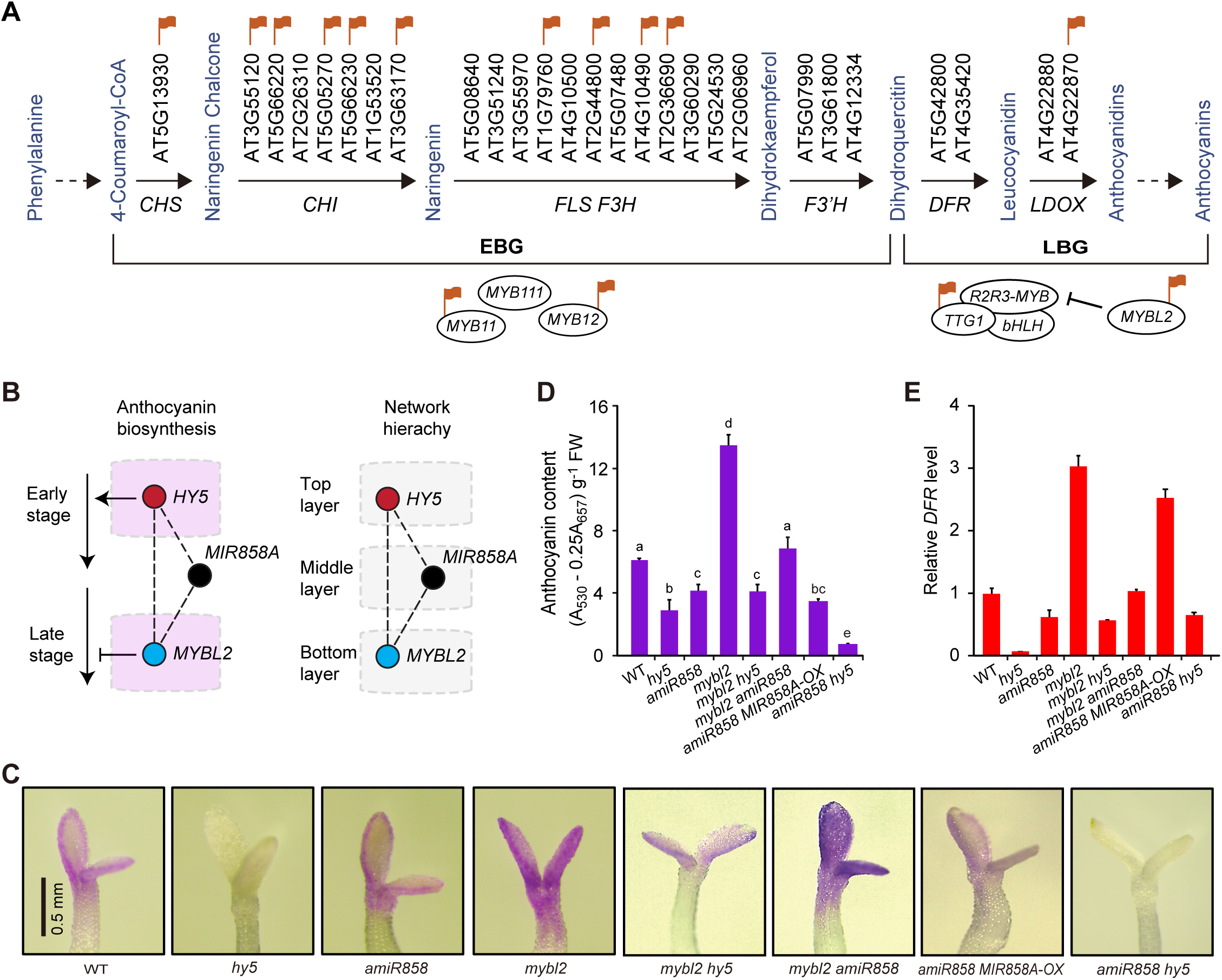
The *HY5-MIR858A-MYBL2* FFL mediates light induced anthocyanin biosynthesis. **A**. Diagram showing the simplified anthocyanin biosynthesis pathway, which involves a series of enzymatic steps divided into the early and late stage. Genes directly bound by HY5 are noted with red flags. Three related MYB proteins, MYB11, MYB12, and MYB111, positively regulate genes encoding early stage enzymes. HY5 binds to the promoter of these genes, except for *MYB111. MYBL2* is pertinent to the late stage due to its interference with the formation of the MBW complex that promotes expression of key genes such as *DFR* and *LDOX*. **B**. Partitioning of the *HY5-MIR858A-MYBL2* FFL in the anthocyanin pathway (left) and the hierarchy of the core network (right). **C**. Visualization of purple pigmentation in the double mutants in comparison to the single mutants. Seedlings were grown in light in the presence of norflurazon and photographed. Bar, 0.5 mm. **D**. Quantitative measurement of anthocyanin content. Data are means ± SD (n = 4). Samples labeled with different letters denote groups with significant differences (one way ANOVA test, *P* < 0.05). **E**. qRT-PCR analysis of mRNA levels of the anthocyanin biosynthetic gene *DFR* in seedlings. All data are means ± SD (n = 3).

To test the relationship between the molecular arrangements of the nodes and their genetic interactions, we employed the *hy5, amiR858*, and *mybl2* single mutants. We generated all three pairwise combinations of these single mutants (*mybl2 hy5, mybl2 amiR858*, and *amiR858 hy5* double mutants). We then assayed light-induced anthocyanin accumulation in these double mutants by direct visualization (Figure 7C), chemical quantification (Figure 7D), and expression profiling of the anthocyanin biosynthetic gene *DFR* (Figure 7E). These analyses revealed that the *mybl2 hy5* and *mybl2 amiR858* mutants exhibited intermediate phenotypes compared to the single mutants (Figures 7C to 7E). Mutations in *HY5* and *MIR858A* appeared to have an additive effect at the physiological level, as *amiR858 hy5* displays lower anthocyanin accumulation than *hy5* (Figures 7C and 7D). Consistent with the molecular arrangements of the three nodes, these results demonstrate that the input node *HY5* has a negative and a positive genetic interaction with *MYBL2* and *MIR858A*, respectively, to coordinately modulate the output node *MYBL2*.

### *MIR858A* contributes to irradiance-dependent pigmentation

To gain further insight into the function of *MIR858A*, we monitored the expression dynamics of the FFL in a light intensity gradient consisting of NL (no light), LL (low light), ML (medium light), and HL (high light). Under these conditions, pigmentation in wild type seedlings progressively increases as the irradiance increases (**Figure 8**A). We found that the *HY5*-overexpressing seedlings (*HY5-OX*) accumulated anthocyanins to higher levels than the wild type, while the *hy5* mutant dramatically abolished light-induced pigmentation across the light gradient (Figure 8A). As expected, gain- and loss-of-function *MYBL2* mutants manifested pigmentation defects opposite to those of the *HY5* mutants across the light gradient (Figure 8A). Conversely, genetic manipulations of *MIR858A* only resulted in drastic phenotypes under HL (Figure 8A).

**Figure 8.**
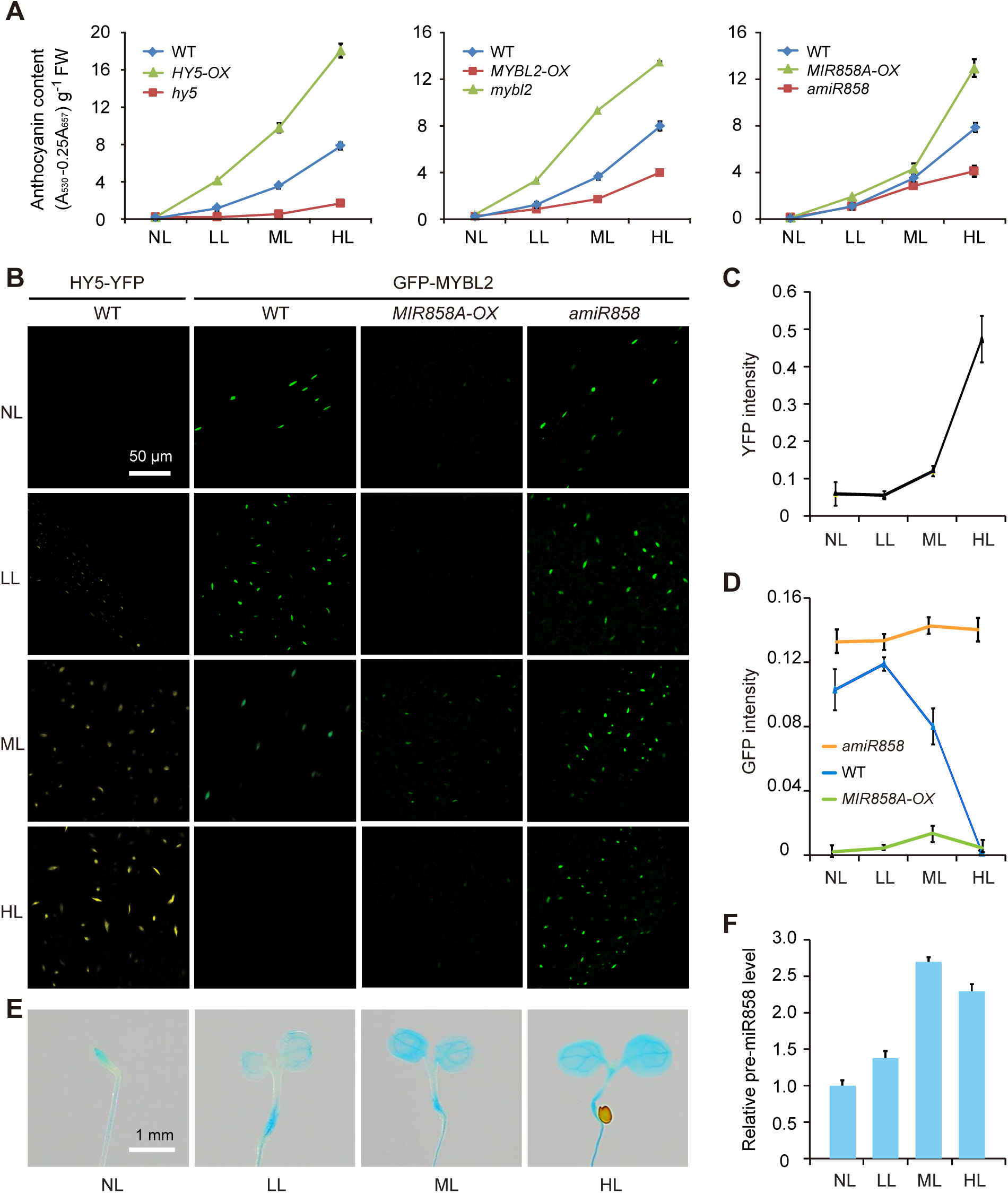
*MIR858A* facilitates the dynamics of the *HY5-MIR858A-MYBL2* FFL in light induced pigmentation. **A**. Quantification of irradiance-dependent anthocyanin levels in *HY5, MYBL2*, and *MIR858A* gain- and loss-of-function mutants in comparison to the wild type. Seedlings were grown for three days in the four indicated light conditions and measured for their total anthocyanin content. Data are means ± SD (n = 4). NL, no light; LL, low light; ML, medium light; HH, high light. **B**. Nuclear accumulation pattern of HY5 and MYBL2 in response to a gradient of light irradiance. Seedlings expressing *HY5-YFP* or *MYBL2-GFP* in the indicated genetic backgrounds were grown under different light conditions. Nuclear-localized fluorescent signals were examined by fluorescence microscopy. Bar, 50 µm. **C**. Quantification of nuclear YFP fluorescence intensity in *HY5-YFP*–expressing seedlings grown under different light conditions. **D**. Quantification of nuclear GFP fluorescence intensity in the wild type, *MIR858A-OX*, and *amiR858* seedlings expressing the same *MYBL2-GFP* reporter grown under different light conditions. Data represent means ± SD (n = 30). **E**. GUS activity detected in seedlings expressing the *pMIR858A:GUS* reporter gene grown under four light conditions. Bar, 1 mm. **F**. qRT-PCR analysis of pre-miR858a transcript levels under the indicated light conditions with values normalized to that of NL. Data are means ± SD (n = 3).

Using a transgenic line expressing HY5 with yellow fluorescent protein (YFP) fused to its C-terminus [42], we observed that the nuclear YFP-HY5 signal progressively increased from NL to HL (Figure 8B). Quantification revealed that the HY5 level increased by approximately 10-fold from NL to HL (Figure 8C), confirming the well-documented phenomenon of light-induced nuclear accumulation of HY5 [43]. For MYBL2, we fused its coding region with green fluorescent protein (GFP) under control of the native *MYBL2* promoter. In plants stably expressing the *pMYBL2:GFP-MYBL2* reporter, the GFP signal markedly decreased from NL to HL (Figure 8B). Quantification of the nuclear fluorescence revealed a >10-fold difference in the level of GFP-MYBL2 between the NL and HL treatments (Figure 8D).

We then introduced *pMYBL2:GFP-MYBL2* into the *MIR858A-OX* and *amiR858* backgrounds. GFP fluorescence increased drastically in *amiR858* but decreased in *MIR858A-OX* seedlings compared to the wild type. At the same time, the light responsiveness of the reporter gene was abolished in both mutant backgrounds (Figures 8B and 8D). Moreover, we profiled the *MIR858A* promoter activity in response to light using the *pMIR858:GUS* reporter line (Figure 8E). This analysis, together with quantification of pre-miR858a levels (Figure 8F), revealed that the *MIR858A* promoter was activated by increasing light intensity. Thus, miR858 level tracked HY5 abundance but was inversely correlated with nuclear MYBL2. However, this activation by light was stronger under ML to HL than in the LL (Figures 8E and 8F). Taken together, the modes of action of the three nodes indicate that *MIR858A* provides a structural basis for facilitating interaction between nodes from different hierarchical layers.

## Discussion

Reconstruction of gene networks integrating multiple regulatory relationships within the appropriate biological context is a desired approach for advancing plant biology [10, 44]. Currently, the reported networks in plants are primarily focused on transcriptional control [7, 15, 45-47]. We mapped and analyzed a directed network in *Arabidopsis* with a focus on the upstream and downstream interactions of miRNAs (Figures 1A and 1B; Figure S1 to S5). Our results highlighted the prevalence of miRNA-containing FFLs in the formation and function of date hubs.

### MiRNAs promote FFL wiring

FFL is a prominent and versatile network motif [9, 10]. In plants, a number of FFLs have been identified in gene regulatory networks and characterized in relation to developmental processes and stress responses [7, 10, 14-16]. In the reconstructed network reported here, the miRNA nodes collectively displayed highly comparable out-degree and in-degree (Figure 1C; Figure S6A). This unique property was associated with a significant enrichment of miRNA-containing FFLs but not topological isomeric FBLs (Figures 2A to 2D). Thus, miRNAs structurally enhance FFL-compatible wiring.

When considering the type of regulation (activating or repressing), there are eight different subtypes of FFLs [9, 11]. Mathematical modeling and experimental monitoring of gene behaviors have demonstrated that different FFL subtypes fulfill distinct functions, such as the detection of expression fold change or protection against premature responses to brief environmental fluctuations [12, 13]. There are two fundamental features of miRNA-containing FFLs. First, in three-quarters of the FFLs, the miRNA was positioned as the intermediate node (Figure 2D and 2E). Second, miRNAs typically repress expression of their target genes. Therefore, the subtype of miRNA-containing FFLs in plants would be determined primarily by the TTI and TMI emanating from the same input TF.

In the exemplar *HY5-MIR858A-MYBL*2 FFL, TTI and TMI from the input node *HY5* had opposite signs of regulation (Figure S8 and S9), establishing the FFL as a coherent subtype (Figure S8A). Consistent with this circuit design, we found that both the direct and indirect path were used to suppress *MYBL2* expression in a light gradient. However, *MYBL2* repression by the input node *HY5* was less effective than the Y node *MIR858A* as manifested by MYBL2-GFP dynamics (Figure 8). Taken together, these data suggest that miRNA-containing FFLs were favorably amplified in *Arabidopsis* through binary topological generalization (Figure 3), taking advantage of the effective post-transcriptional repression mechanism to achieve quantitative silencing of multiple targets. Through this binary topological generalization mechanism, certain miRNAs are shaped into party or date hubs connected as intertwined FFLs (Figure 4 and 5).

### Choreographing of light-induced pigmentation by a miRNA-containing FFL

As an illustrative example, we elucidated *HY5*-*MIR858A*-*MYBL2* as a decision-making switch controlling light-induced pigmentation (Figure 7), extending previous knowledge regarding its individual nodes [34, 35, 40, 41, 49]. Light perception and activation of the loop is accomplished via HY5 accumulation in the nucleus [43, 50]. Consistently, we found that constitutive accumulation of HY5, which is a top-layer node in the core network, was sufficient to activate the anthocyanin biosynthesis pathway (Figures 7 and 8). Genetic analysis further demonstrated that *MIR858A* was indispensable for this programming by quantitatively responding to *HY5* and repressing *MYBL2* expression (Figures 7 and 8), establishing a reciprocal nuclear accumulation pattern of the X and Z nodes in response to a gradient of light irradiance (Figure 8).

Together with known molecular mechanisms of the light signaling and anthocyanin biosynthesis modules, our findings support a “seesaw” model for explaining light intensity dependent pigmentation in *Arabidopsis* seedlings via the *HY5*-*MIR858A*-*MYBL2* FFL. In darkness, the balance of the two antagonist TFs is tipped toward the low-HY5-high-MYBL2 state due to proteolytic removal of HY5 by CONSTITUTIVELY PHOTOMORPHOGENIC 1 (COP1), which encodes a RING-finger E3 ubiquitin ligase [43, 51]. Because HY5 and MYBL2 act mainly to promote early genes and inhibit late genes, respectively (Figure 7A), this low-HY5-high-MYBL2 state shuts down anthocyanin biosynthesis. An advantage of this design is that brief fluctuations in light irradiance may be ignored, as it takes time for miR858 to reach sufficient levels to repress MYBL2. The FFL is thus a possible fail-safe mechanism to reliably prevent pigmentation without prolonged light irradiance.

Upon exposure to continuing light, COP1 is rapidly reduced to allow nuclear accumulation of HY5 [43, 50]. For anthocyanin biosynthesis to take place, however, the direct path of HY5-mediated transcriptional suppression of *MYBL2* is not sufficient without miR858 (Figure 7). Inhibition of MYBL2 accumulation in the nucleus is mainly outsourced to *HY5*-activated *MIR858A*, effectively converting the light signal into a high-HY5-low-MYBL2 state for active anthocyanin biosynthesis. Taken together, we showed that the *HY5*-*MIR858A*-*MYBL2* FFL is a decision-making module that equips the plant with the ability to quantitatively interpret light input and tip the “seesaw” of anthocyanin production accordingly, resulting in pigmentation proportional to the perceived irradiance (Figure 5 and Figure S10).

### Global relationship between TFs and miRNAs

In both animals and plants, miRNAs are known to have a higher propensity to interact with TFs [1, 2, 22]. The in-degree distribution of animal miRNA networks has been shown to follow a power law, and so-called target hubs acted upon by multiple miRNAs have been identified [52]. Furthermore, these target hubs are enriched with TFs in diverse species [52], indicating their evolutionary conservation in the animal lineage. No predicted or validated miRNA-targeted hubs have been reported in plants. Consistently, we found no evidence supporting the existence of miRNA target hubs in the MTIs, possibly owing to the high degree of complementarity required for sufficient miRNA action in plants [2, 22].

In contrast to the prevalence of intronic miRNAs in animals [53], most plant miRNA loci are encoded as independent transcription units [8, 24, 25]. TMIs are therefore important in specifying the spatiotemporal expression domains of miRNAs in plants. In these contexts, we believe the miRNA-containing FFLs have two implications in the global relationship between TFs and miRNAs. First, with their non-miRNA nodes strategically positioned in different hierarchical layers (Figure 6), miRNA-containing FFLs facilitate relay of regulatory information from the input node to the output node (Figure 8). This function may reinforce the “vertical” hierarchy and thus could be advantageous in fixing a miRNA node along with its silencing interactions in the network (Figure 7). Second, the vertical hierarchy may have an “oriented” binary topological generalization that arrived at either an input- or an output-heavy structure. Taken together, we speculate that the party and date miRNA hubs were derived from expanding miRNA-containing FFLs, because these miRNAs were effective in carrying out the Y node function. As a result, the selected FFLs facilitate “horizontal” crosstalk among different functional modules [26]. Thus, the overall architecture of TF-miRNA regulatory networks appears to be different in animals and plants, although further systems level analyses are required to form definite conclusions.

In summary, we found that miRNAs occupy a distinct niche than other nodes in the reconstructed miRNA network of *Arabidopsis*. This finding shed new light on the global role that miRNAs play in shaping the architecture and organization of gene networks. While still incomplete, the “wiring diagram” connecting miRNAs with other genes represents a useful framework for understanding combined transcriptional and post-transcriptional regulatory mechanisms. This, together with the ability to genetically manipulate miRNAs and assess the transcriptomic and phenotypic consequences, will allow the design principles and control logics of gene expression programs to be deciphered with increasing detail and clarity [10].

## Materials and methods

### Plant materials and growth conditions

*Arabidopsis thaliana* ecotype Col-0 was used as the wild type plant for all experiments. Mutants defective in *HY5* and *BZR1* were *hy5-215* [49, 54] and *bzr1-D* [38], respectively. For MYB63 and *MYBL2*, the T-DNA insertion lines *SALK_049267, SALK_092920C, SALK_107780*, and *SALK_126807* were used. The *35S:HY5-YFP* line used was as previously described [42]. To constitutively activate *HY5*, the coding region together with the 3’ UTR driven by the *Ubiquitin-10* promoter was cloned into the pCAMBIA1300 vector (CAMBIA) using the primers listed in Table S2. The construct was introduced into wild type plants by standard *Agrobacterium* mediated transformation. Transformants were selected by Hygromycin resistance, and T_2_ progenies were used in subsequent experiments. For *MYBL2* overexpression, the coding region was inserted into the pCAMBIA1305.1 vector downstream of the CaMV *35S* promoter. T_3_ generation seedlings were used for phenotypic analysis.

For overexpression of *MIR858A*, the genomic sequence encompassing pre-miR858a was amplified using the primers listed in Table S2. The artificial miRNA *amiR858* was generated using the pre-miR319 backbone and bridge PCR with primers listed in Table S2. The PCR products were inserted into the pJim19 vector under control of the CaMV *35S* promoter. The two constructs were introduced into wild type plants and selected with Basta resistance and allowed to propagate to the T_3_ generation. To generate the *amiR858 MIR858A-OX, mybl2 hy5*, and *mybl2 amiR858* double mutants, F_2_ plants homozygous for both alleles were selected with appropriate antibiotics followed by PCR analysis of genomic DNA. F_3_ progenies were used in subsequent experiments.

To create GUS reporter lines, the 1332 bp and 1933 bp genomic regions upstream of pre-miR858a and the start codon of *MYBL2* were cloned. The *pMIR858A* sequence was first cloned into the TOPO vector (Invitrogen). Deletion of the two core sequences (ACGT) of the distal and proximal G-box individually and together was achieved through PCR using the primers listed in Table S2. The four resulting constructs were then subcloned into the pCAMBIA1381-Xa vector. The *pMYBL2:GUS* reporter gene was cloned into the pCAMBIA-1305.1 vector. These constructs were used to transform wild type plants. The *pMIR858A:GUS* and *pMYBL2:GUS* reporter genes were also introduced into the *hy5* background through genetic crossing. Homozygotes were selected from seedlings that exhibited long hypocotyls and resistance to the appropriate antibiotics. To generate the *pMYBL2:GFP-MYBL2* construct, cloned DNA fragments corresponding to the 1933 bp *pMYBL2*, the 729 bp coding region of GFP, and the 588 bp coding region of *MYBL2* were sequentially inserted into the pCAMBIA1300 vector. The resulting contruct was used to transform wild type plants and subsequently introduced into *MIR858A-OX* and *amiR858* through genetic crossing. F_1_ seedlings were screened for resistance to both Hygromycin and Basta and monitored for GFP fluorescence. This process was repeated for the F_2_ progenies.

To grow *Arabidopsis* seedlings, seeds were surface sterilized and plated on agar-solidified MS media including 1% (w/v) sucrose. The plates were incubated at 4°C for 3 days in the dark and then transferred to a growth chamber with a 22°C/20°C, 16 h light/8 h darkness setting. Adult plants were maintained in standard long-day (16 h light/8 h darkness) conditions, with a light intensity of 120 μmolm^−2^s^−1^, 50% relative humidity, and a temperature of 22°C. Tobacco plants (*Nicotiana benthamiana*) were maintained under the same conditions, except for the temperature was set at 25°C and light intensity at 200 μmolm^−2^s^−1^.

### MiRNA target identification

Sequences of the 428 *Arabidopsis* miRNAs were obtained from miRBase (version 22) [55]. Computational prediction using psRNATarget [31] and psRobot [30] was based on two filtering scores: 2.2 or 2.5 (default) in psRNATarget and 2.5 or 3.0 (default) in psRobot. The standard for filtering the results was the penalty score for the miRNA-target alignment. The four outputs were then searched against degradome sequencing data processed by the CLEAVBELAND pipeline [56, 57]. Possible MTIs predicted by both programs or by either program but compatible with the degradome data were combined into four datasets and tested against a benchmark consisting of 449 validated miRNA targets. Output from the psRNATarget (2.5) and psRobot (2.5) combination was selected to represent MTIs because it offered a suitable tradeoff between coverage and false positive rate (Figure S1). Association of the target genes with GO terms was analyzed using AgriGO. Fisher’s exact test and the Yekutieli [false discovery rate (FDR) under dependency] method were used to detect enriched terms with FDR set at 0.05.

### ChIP data processing and annotation

All ChIP data were obtained from published works. For ChIP-seq data, the raw data in the SRA storage mode was converted to fastq format and sequentially quality checked using fastQC and Cutadapt [58] to remove remaining adaptors, overrepresented false fragments, low quality reads, and unrecognized nucleotides (marked with N). The clean data was mapped to the TAIR10 reference genome using Bowtie [59] allowing no more than one mismatch, and only the best-mapped site for each read was retained. Samples with a low mapping rate (< 50%) were discarded. TF-binding peaks were called using MACS [60] with default parameters except that the genome size was set as -g (1.1e^8^) and FDR typically as -p (0.001). Biological replicates were treated as previously described [45]. Briefly, for replicates with significantly different mapping rates, only samples with the highest mapping rate were retained. If the replicates had similar mapping rates, their Pearson correlation coefficient was calculated and the replicate with the highest value was retained.

For ChIP-chip data, the probes were mapped to the TAIR10 genome build using STAR [61]. Raw CEL data files were normalized and analyzed using TAS (Affymetrix tiling array software). The TAS software was obtained from the Affymetrix website (https://www.thermofisher.com/cn/zh/home.html). Probe intensity was computed based on both PerfectMatch and MisMatch (PM/MM) with a bandwidth of 300 bp. The binding peaks were defined as those with *P* < 0.05 with maximum gap of 300 bp and minimum run of 100 bp. The peaks were detected using the TileMap Peak Detection suite with MA set as the Region Summary Method, half window size as five probes and 125 bp, and other parameters as default.

Custom Perl scripts were used to assign the identified peaks to genomic loci following the principles described previously [62, 63]. For each protein-coding locus, a window extending from 2000 bp upstream to 300 bp downstream of the TSS was defined. For a miRNA locus, the window was defined as 2000 bp upstream of the first base in the pre-miRNA. A peak was assigned to a gene if the midpoint of the peak fell within the given window.

### Network reconstruction and analysis

The network incorporating TTIs, TMIs, and MTIs was reconstructed using Cytoscape (version 3.4.0) with the Edge-weighted Spring Embedded Layout method [64]. Global topology was examined using the NetworkAnalyzer module [64]. Mfinder software [65] was used to compute the enrichment of network motifs with no greater than four nodes. Permutation that preserved the number of nodes and edges but randomized the connections was repeated 1000 times. The Pearson correlation coefficient of the fragments per kilobase per million reads (FPKM) for peak regions was calculated using the R function cor.test. Statistical analyses were performed using the R function chisq.test and t.test, which were based on independent-samples t-test (t.test(x,y,paired=FALSE)) after checking the data distribution by R library (car).

The TF-miRNA core network was analyzed using the hierarchical layout in Cytoscape. The nodes were divided into three layers based on degree level and clustering coefficient in comparison to the calculated average clustering coefficient. Specifically, nodes with a clustering coefficient greater than the average clustering coefficient were grouped as one layer. Nodes with a clustering coefficient less than the average clustering coefficient were further divided into two layers (those with in-degree less than 10 but an out-degree no less than 10 and those with an in-degree no less than 10 but an out-degree less than 10). Relative expression levels for these nodes were determined using RNA-seq data. The quality of the data was checked by fastQC with adapter sequences removed and low quality bases further trimmed using Cutadapt. Reads were mapped to TAIR 10 using STAR. Cuffdiff [66] was employed for differential expression analysis.

To analyze hub miRNAs, the degree level and clustering coefficient for all miRNA nodes were determined by Cytoscape. Four types of hub miRNAs were identified. Criterion for defining in hubs and out hubs was in-degree greater than ten and an out-degree greater than ten, respectively. Criteria for defining party hubs and date hubs were an out-degree greater than five and a clustering coefficient greater than the network average. These two types of hubs were further classified based on manual curation of GO terms associated with the target genes. Those hubs with targets associated with similar GO terms were classified as party hubs, and those with targets associated with dissimilar GO terms were defined as date hubs.

### ChIP-qPCR

Chromatin isolation was performed using 4-day-old whole seedlings grown under HL. The chromatin pellet was resuspended and sonicated at 4°C to a mean size of approximately 500 bp using a Bioruptor (Diagenode). Sheared DNA was immunoprecipitated with a polyclonal HY5 antibody purified from anti-HY5 rabbit IgG as previously described [7]. Flow-through IgG without HY5 antibody was used as control. Along with an aliquot of sonicated DNA without further treatment (input), the samples were washed, reverse cross-linked, and subjected to qPCR analysis. PCR products from the HY5 antibody and IgG treated samples were normalized against those from the input DNA, and the fold of enrichment was calculated.

### Quantitative transcript analysis

Total RNA was isolated using Trizol reagent (Catalog No. 15596018, Thermo Fisher) and was treated with DNase I (Catalog No. 2270A, Takara, Kusatsu, Shiga) according to the manufacturer’s instructions. RNA was treated and reverse transcribed using the PrimeScript™ II 1^st^ Strand cDNA Synthesis Kit (Catalog No. 6210A, Takara, Kusatsu, Shiga) according to the manufacturer’s instructions. qPCR was performed with SYBR Green master mix on the ABI 7500 Fast Real-Time PCR System (Applied Biosystems). The *Actin7* gene was used as an internal control and normalization standard. Quantification of miRNA levels was carried out using the miRcute miRNA Isolation Kit (Catalog No. DP501, Tiangen, Beijing) for isolation of low-molecular-weight RNA, the miRcute miRNA First-Strand cDNA Synthesis Kit (Catalog No. KR211-01, Tiangen, Beijing) for poly(A) tailing and first-strand cDNA synthesis, and the miRcute miRNA qPCR Detection Kit (Catalog No. FP411-02, Tiangen, Beijing) for qPCR analysis. 5S ribosomal RNA was used as an internal control. Each qPCR experiment included three independent biological replicates and was repeated at least three times. Data from one representative experiment were shown in the text.

### Assay for GUS activity

Seedlings expressing various *GUS* reporter constructs were grown for 3 days in HL and then subjected to various light treatments as indicated. Harvested seedlings were immersed in GUS staining solution containing 1 mM X-Gluc (Catalog No. 18656-96-7, INALCO, San Luis Obispo, California) for 3 h at 37°C as described [67]. Following removal of the staining solution, chlorophyll was washed away with 75% ethanol. Images of the seedlings were acquired with a digital camera.

### Anthocyanin assays

Anthocyanin visualization in seedlings was facilitated by treatment with 100 μM Norflurazon (Catalog No. 34364, Sigma-Aldrich) as previously described [40]. Images of 3-day-old seedlings grown under various light conditions were documented with a stereomicroscope equipped with a digital camera (Leica). For quantification of anthocyanin content, 3-day-old seedlings grown on MS medium were harvested and homogenized. Pigment extraction and determination of anthocyanin levels were performed as previously described [68].

### Dual luciferase assay

The *35S:pre-miR858a* effector construct was generated based on the pGreenII 62-SK vector using the primers listed in Table S2. The *pACT2:MYBL2-LUC* reporter construct was generated using the pGreen II 0800-LUC vector with the 1499 bp *Arabidopsis ACTIN2* promoter cloned in front of the *MYBL2* coding sequence with the TGA stop codon removed. This construct was further modified through three rounds of PCR to mutate the nucleotide sequence of the MBS without changing the amino acid sequence to generate *pACT2:MYBL2*^*mMBS*^*-LUC*. The *pMYBL2:LUC* construct was generated by inserting the *MYBL2* promoter into pGreen II 0800-LUC. Protoplast isolation from leaves of 4-week-old tobacco plants and DNA transfection were performed following established protocols. Briefly, protoplasts were transfected with paired effectors and reporters (7 μg of DNA per construct) and incubated for 16 h in the dark. Transformed protoplasts were collected, homogenized, and dual luciferase reactions carried out using the Dual-Glo Luciferase Assay System (Catalog No. E1501, Promega, Madison). Luciferase activity was quantified using a Multimode Reader LB 942 luminometer (Berthold).

### Measurement of lignin content

The main inflorescence stem from various genotypes was collected according to a temporal progression scheme delineating inflorescence development [69]. Stem samples were ground in liquid nitrogen, lyophilized, and used to prepare extractive-free cell wall residues as previously described [70]. Lignin content was determined using the acetyl bromide method as described [71].

### Fluorescence microscopy

*Agrobacterium* GV3101 cells harboring the *pMYBL2:GFP-MYBL2, SV40:mCherry* [72][71], and *Tav2b* constructs were mixed in a 6:1:2 ratio and used to infiltrate young tobacco leaves. Images were acquired 3 days later with a LSM710 laser scanning confocal microscope (Zeiss), using 488 nm excitation, 490-560 nm emission wavelength for GFP and 543 nm excitation, 575-797 nm emission wavelength for mCherry. For YFP and GFP fusion proteins stably expressed in *Arabidopsis*, confocal microscopy was carried out in a similar manner. Measurement of GFP and YFP fluorescence intensity was performed using ImageJ software. Lignin autofluorescence was excited at ∼400 nm and collected at 420-560 nm.

## Data availability

Sequence data from this article can be found in the *Arabidopsis* Genome Initiative or GenBank/EMBL databases under the following accession numbers: *MIR858A* (At1g71002), *HY5* (At5g11260), and *MYBL2* (At1g71030). ChIP sequencing data are deposited under GSE45938, GSE20176, GSE46986, GSE21301, GSE38358, GSE53422, GSE48793, GSE30711, GSE48082, GSE51120, GSE51537, GSE48081, GSE24568, GSE38358, GSE39215, GSE35315, GSE35059, GSE36361, GSE49282, GSE26722, GSE14600, GSE46986, GSE45846, GSE45213, GSE33120, GSE35952, GSE56706, GSE43637, GSE70533, GSE68193, GSE60084, GSE60554, GSE71397, GSE59187, GSE63463, GSE69431, GSE66290, GSE76571, GSE64245, GSE16940, SRP005412, SRP017902, and GSE80568. ChIP microarray data are deposited under GSE17717, E-MEXP-2653, GSE44872, GSE24684, GSE19763, GSE43291, GSE36965, GSE13090, GSE13090, GSE40519, GSE24974, GSE28063, GSE14635, E-MEXP-2068, GSE33297, GSE33297, and E-MEXP-2499. RNA sequencing data are deposited under GSE77211, GSE80712, and GSE52407.

## Authors’ contributions

LL (Lei Li) designed the research. ZG, HH, and LL (Lei Li) analyzed the data. ZG collected the data and performed the bioinformatics analyses. JL (Jun Li), YY, JL (Jian Li), CF, DZ, and HC performed the research. JL (Jun Li) performed experiments. YY quantified the pre-miR858 level. JL (Jian Li) and DZ provided *HY5-OX, 35S:HY5-YFP*, and *bzr1-D* seeds. CF quantified the lignin contents. HC constructed the *pMIR858A:GUS, pMIR858A*_*m1*_:*GUS, pMIR858A*_*m2*_:*GUS*, and *pMIR858A*_*m1+2*_:*GUS* vectors. ZG, JL (Jun Li), and LL (Lei Li) wrote the paper. ZG and LL (Lei Li) revised the paper. All authors approved the final manuscript.

## Acknowledgments

This work was supported by the National Key Research and Development Program of China (2017YFA0503800) and the National Natural Science Foundation of China (31621001).

## Supplementary material

**Supplemental Figure S1 Compilation of MTIs through computational prediction and degradome sequencing analysis**

**A**. Generation of four datasets of putative MTIs based on computational predictions. Outputs from psRNATarget (maximum expectation score set as 2.5 or 3.0) and psRobot (penalty score set as 2.2 or 2.5) were compared against degradome sequencing data. Four subsequent datasets were generated with each, including putative targets predicted by both programs or by either program but compatible with degradome data. **B**. Selection of the optimal dataset for representation of MTIs. The four datasets were tested against a benchmark of 449 validated targets to calculate recovery rate and total number of predictions. The dataset that combined output from psRNATarget (2.5), psRobot (2.5), and the degradome data was considered optimal. **C**. Pie graph showing the final set of 2823 MTIs that includes 2712 predictions and 111 canonical targets not recovered by prediction. **D**. Distribution of the MBS along the target genes. The 2008 target genes from the 2823 MTI were aligned from head to tail and divided into 20 intervals. The number of MBSs in each interval was calculated and plotted.

**Supplemental Figure S2 TFs are enriched in miRNA target genes**

**A**. TF families targeted by miRNAs. A total of 35 TF families in *Arabidopsis* were found to include at least one member targeted by MTIs. **B**. Enrichment of TFs in miRNA targets. The proportion of TFs targeted by miRNAs was significantly higher than that at the genome level. *** represents *P* < 0.001 by chi-square test. **C**. Bubble graph showing GO term analysis of miRNA target genes. The horizontal coordinate represents the factor of enrichment, which is the relative frequency for a given term in the query set against that in the genome. The vertical coordinate represents enriched GO terms in the miRNA targets, with a false-discovery rate (FDR) cutoff of 0.05. The size of the ovals represents the number of genes associated with a given term. The colors of the ovals represent the Q values that indicate the minimum FDR at which significant enrichment of a GO term was determined.

**Supplemental Figure S3 Distribution patterns of TF-binding peaks within the genome**

**A**. The 66 core TFs with qualified global ChIP data were divided into eight non-overlapping groups based on their annotated functions. **B**. Chromosome-level distribution of TF binding peaks. A total of 339,875 binding peaks were identified for the 66 TFs after quality control and uniform processing. Each of the five chromosomes was divided into 1000-bp bins, and a sliding window was used to calculate the number of binding peaks. The average number of binding peaks in each bin was plotted against the chromosomal coordinates. **C**. Localization of the binding peaks in relation to annotated genome components. Percentages of the binding peaks located in 3’ UTR, 5’ UTR, intronic, exonic, and intergenic regions are shown.

**Supplemental Figure S4 Mapping TTIs and TMIs using the TF binding peaks**

**A**. Gene-level distribution of TF binding peaks for protein-coding loci. A sliding window analysis showed that binding peaks concentrate at the TSS and extend 2000 bp upstream and 300 bp downstream of the TSS. This region was defined as a binding window. If the midpoint of any of the 339,875 binding peaks was identified in the binding windows, a putative TTI was declared between the corresponding TF and the given protein-coding gene. **B**. For miRNA loci, the binding peaks were found to concentrate at the 200 bp region upstream of the first nucleotide of the annotated pre-miRNAs. The 2000 bp region upstream of the first nucleotide of the pre-miRNA was defined as the binding window for establishing TMIs for the miRNA loci.

**Supplemental Figure S5 Topological analysis of the reconstructed miRNA network**

**A**. Distribution of node degree in the network. The degree of all nodes was calculated and the frequency distribution plotted, which followed a power-law. **B**. Distribution of the node in-degree. Only the in-degree of all nodes was calculated and plotted. **C**. Histogram showing frequency distribution of the path length. All connected nodes in the network were used to extract the paths, and their lengths (steps) were calculated. **D**. Node properties in the network. Using the NetworkAnalyzer function in Cytoscape, clustering coefficient, closeness centrality, and betweenness centrality for all nodes in the network were determined and their respective distributions plotted. **E**. Distribution of clustering coefficient, closeness centrality, and betweenness centrality after all miRNA nodes were removed.

**Supplemental Figure S6 Enrichment of miRNA-containing network motifs in the reconstructed miRNA network**

**A**. Diagram showing the in-degree (regulated edges) and out-degree (regulating edges) of a given node. **B**. Enriched miRNA-containing motifs (Z score > 2) in the reconstructed network. Graphic representation of the motifs and a specific example for each enrich motif are shown.

**Supplemental Figure S7 The miRNA network contains eight modular TF-defined sub-networks**

**A**. Bubble chart displaying the number of nodes and edges of the eight TF-defined sub-networks together with the pan-network. The size of the disks represents the number of edges. **B**. Visualization of the eight sub-networks reconstructed from the pan-network based on the distinctive functions of the core TFs. The core TFs are shown as colored triangles, miRNAs as black circles, and other genes as grey squares. All edges are shown as grey lines. **C**. Node clustering coefficient at the sub-network and individual TF levels. Values are the average clustering coefficient for the eight sub-networks and the average of all nodes connected to a TF through TTIs and TMIs.

**Supplemental Figure S8 Validation of *HY5-MIR858A-MYBL2* as a coherent FFL**

**A**. Structure of the *HY5-MIR858A-MYBL2* FFL, which consists of three edges forming a direct path and an indirect path. All three regulatory interactions were validated by multiple experimental approaches, which collectively manifested the FFL as a coherent type. **B**. HY5 occupancy at the *MIR858A* locus based on global ChIP data mapped onto genome coordinates. The pre-miR858a and promoter region are depicted as a black arrow and a horizontal bar, respectively. The triangles mark the two G-boxes in the promoter region. P1 and P2 indicate the positions of the two amplicons for ChIP-qPCR analysis. **C**. Confirmation of HY5 binding to the *MIR858A* promoter by ChIP-qPCR analysis. ChIP was performed in the wild type and *hy5* seedlings using an anti-HY5 antibody. The precipitated DNA was analyzed by qPCR, with the values normalized to those of the IgG-treated samples. **D**. Analysis of mature miR858 transcript levels under light and dark conditions. Levels of mature miR858 determined by qRT-PCR were normalized, and the value for light-grown wild type seedlings was set to 1. Data for ChIP-qPCR and qRT-PCR represent means ± SD (n = 3). **E**. GUS activity driven by various forms of the *MIR858A* promoter expressed in the wild type or *hy5* background. Seedlings grown under dark (top) and light (bottom) conditions were stained for GUS activity and visualized. Bar, 1 mm.

**Supplemental Figure S9 *HY5* partially mediates suppression of *MYBL2* transcription by light**

**A**. qRT-PCR analysis of *MYBL2* transcript levels in wild type and *hy5* seedlings grown in the dark and light. *MYBL2* levels were normalized, and the value for wild type in light was set to 1. Data are means ± SD (n = 3). **B**. Analysis of HY5 dependent *MYBL2* promoter activity. Transgenic seedlings expressing *pMYBL2:GUS* in the wild type or *hy5* background were subjected to different light treatments and assayed for GUS activity. Dark-light, transitioning from dark to light for 12 h. Note the difference in seedling morphology influenced by the light treatments. Bar, 1 mm. **C**. Suppression of *MYBL2* promoter activity by HY5. The REN/LUC dual luciferase reporter construct, in which *LUC* is driven by the native *MYBL2* promoter, was employed. Tobacco protoplasts were transformed with this reporter alone or together with *35S:HA-HY5* and incubated overnight in either the dark or light. The ratio of the LUC/REN chemiluminescence values was then determined. Data are means ± SD (n = 2).

**Supplemental Figure S10 Molecular analyses of intertwined FFLs centered on *MIR858A***

**A**. Promoter activity of *MIR858A* and *MYBL2* in response to eBL treatments. Seedlings expressing *pMIR858A:GUS* (top) or *pMYBL2:GUS* (bottom) were treated with ethanol or eBL and stained for GUS activity. Bar, 1 mm. **B** and **C**. Analysis of pre-miR858a and mature miR858 transcript levels in seedlings after eBL (B) and PPZ treatment (C) in comparison to the respective mock treatment by qRT-PCR analysis. **D**. Change in *MYBL2* transcript level in seedlings following eBL and PPZ treatments. **E**. Exogenous eBL and PPZ treatment drastically increases and decreases anthocyanin levels, respectively, in *Arabidopsis* seedlings. Samples labeled with different letters denote groups with significant differences (one way ANOVA test, *P* < 0.05). All eBL and PPZ treatments were 0.1 µM. **F**. Comparison of *MYB63* transcript levels in the inflorescence stem of the *hy5* and *amiR858* mutants to that of the wild type. All qRT-PCR data are means ± SD (n = 3).

**Supplemental Figure S11 Analysis of TFs related to the core network**

**A**. Heatmap displaying the relative expression levels of the 1717 annotated TFs in *Arabidopsis* mutants with defective small RNA pathways. Log_2_-transformed expression levels in *rdr6* and *ago1* mutants compared to that of the wild type are shown. **B**. Analysis of the 249 TFs in the miRNA-TF core network. The proportions of TFs that were miRNA targets and non-targets are shown as stacked bars on the left. The pie graphs in the middle and on the right show family information for the miRNA-targeted TFs.

